# Structure of the Femoral Chordotonal Organ in the Oleander hawkmoth, *Daphnis nerii*

**DOI:** 10.1101/2024.07.10.602854

**Authors:** Simran Virdi, Sanjay P. Sane

**Affiliations:** National Centre for Biological Sciences, Tata Institute of Fundamental Research, GKVK Campus, Bellary Road, Bengaluru, India

## Abstract

Insect legs serve as crucial organs for locomotion and also act as sensory probes into the environment. They are involved in several complex movements including walking, jumping, prey capture, manipulation of objects, and self-grooming. These behaviours require continuous modulation of motor output through mechanosensory feedback which is provided by numerous mechanosensors located on the cuticle and within the soft tissue. A key mechanosensory organ in the insect leg, the femoral chordotonal organ (FeCO), detects movements of the femoro-tibial joint. This organ is multifunctional and senses both self-generated movements (proprioception) and external stimuli (exteroception). Movements of the tibia alter the length of FeCO, which activates the embedded mechanosensory neurons. Due to the mechanical nature of these stimuli, the structure and material properties of the FeCO are crucial for their function, alongside neural encoding properties. Here, as a first step towards understanding how its structure modulates its function, we characterized the morphology and anatomy of FeCO in the hawkmoth *Daphnis nerii*. Using a combination of computed micro-tomography, neuronal dye fills and confocal microscopy, we describe the structure of FeCO and the location, composition and central projections of FeCO neurons. FeCO is located in the proximal half of the femur and is composed of the ventral (vFeCO) and dorsal scoloparia (dFeCO) which vary vastly in their sizes and the number of neurons they house. The arrangement of neurons within dFeCO follows a decreasing size gradient in the proximo-distal axis. The characteristic accessory structures of chordotonal organs, the scolopales, significantly differ in their sizes when compared between the two scoloparia. FeCO neurons project to the central nervous system and terminate in the respective hemiganglia. Using these morphological data, we propose a mechanical model of FeCO, which can help us understand several FeCO properties relating to its physiological function.

## 1. Introduction

Insect legs exhibit a vast array of forms and functions, mirroring the remarkable diversity found among insects. Beyond their role in locomotion, insect legs also serve as highly sensitive sensory organs. They perform complex tasks such as walking (Hughes, 1952), jumping (Brown, 1967), landing on substrates (Chapman, 1998; Goodman, 1960), prey capture (Maldonado et al., 1967), swimming and self-grooming (Chapman, 1998) etc, which are actively driven by motor commands that depend on sensory feedback. The fine motor control required for these tasks is provided by multiple mechanosensors in the legs which offer both proprioceptive input during body positioning and self-movement, and exteroceptive input including wind and touch (Delcomyn et al., 1996). Some mechanosensors such as campaniform sensillae are present on the cuticle and sense strain patterns due to load changes in the exoskeleton (Pringle, 1938a; Spinola & Chapman, 1975). On the other hand, hair sensillae (or mechanosensory bristles) sense tactile input from external agents and inter-segmental movements (Pringle, 1938b; Wong & Pearson, 1976). In contrast, chordotonal organs are internally located within the soft tissue. They function as stretch receptors, spanning jointed segments to sense the positions and movements of the distal segment relative to the proximal segment that houses them (Field & Matheson, 1998). Chordotonal organs may be connected to the exoskeleton (connective) and function as proprioceptors, or to internal tissues like the trachea (non-connective) and primarily play a role in audition (Field & Matheson, 1998).

Among the various chordotonal organs within an insect’s body, the femoral chordotonal organ (FeCO) is a crucial mechanosensory organ for many leg-related functions. It is located in the femur and monitors the femoro-tibial joint of each leg. Composed of several closely packed neurons, FeCO encodes several aspects of the femoro-tibial (FT) joint state including the angular position of tibia relative to femur, the speed and direction of the tibial movement, and the substrate vibrations transmitted *via* the tarsi and tibia (Burns, 1974; A. Büschges, 1994; Field & Pflüger, 1989; Hofmann et al., 1985; Mamiya et al., 2018; Matheson, 1990; Stein & Sauer, 1999; Theophilidis, 1986; Usherwood et al., 1968; Zill, 1985).

Studies in locusts (Shelton et al., 1992), stick insects (Hofmann et al., 1985), crickets (Nowel et al., 1995), moths (Kent & Griffin, 1990) and flies (Mamiya et al., 2023) demonstrate that the FeCO is mechanically coupled to tibia *via* a rigid rod-like structure called the *FeCO receptor apodeme*. Apodemes are invaginations of the exoskeleton which serve as attachment points for connective chordotonal organs (receptor apodemes) and muscles (muscle apodemes). The FeCO apodeme originates from the tibial cuticle and projects into the femur where it attaches to the organ *via* a ligament (Field & Matheson, 1998; Nowel et al., 1995; Shelton et al., 1992). Thus, flexing the FT joint causes the apodeme to move distally, stretching the FeCO and extension of the joint moves the apodeme proximally, thereby shortening the FeCO (Field & Matheson, 1998). It thus transmits the rotational stimulus from the tibia to the FeCO in the form of stretch and relaxation.

FeCO is composed of 1-3 subunits called *scoloparia*. This morphological feature is conserved across insects and may be essential for function. For instance, the proximal and distal scoloparia in the locust pro– and meso-thoracic FeCO are vibration-sensitive and movement-sensitive respectively and have distinct central projection patterns (Field & Pflüger, 1989). The FeCO of the vinegar fly *Drosophila melanogaster*, a holometabolous insect, has three scoloparia (Shanbhag et al., 1992) with one scoloparia housing the vibration sensitive *club* neurons while the other two house a combination of position sensitive *claw* neurons and direction sensitive *hook* neurons (Mamiya et al., 2018, 2023). The three functional classes also have distinct central projection patterns from which they derive their names (Mamiya et al., 2018; Phillis et al., 1996). FeCO scoloparia thus exhibit functional specialization across insects.

Within FeCO scoloparia, the neurons are divided based on their sensitivity to position, speed, direction and vibration frequencies. In locusts, the overall activity of the FeCO and its position-sensitive units increases towards the extended and flexed joint angles (Burns, 1974; Usherwood et al., 1968; Zill, 1985) and shows hysteresis upon cyclic stimulation (Burns, 1974; Matheson, 1990; Zill, 1985). The velocity-sensitive units respond to either unidirectional or bidirectional movement, in combination with certain positions in locusts and stick insects (Büschges, 1994; Hofmann et al., 1985; Matheson, 1990). A subset of FeCO neurons are sensitive to low-amplitude, high-frequency vibrations of the FT joint in locusts (Field & Pflüger, 1989), stick insects (Stein & Sauer, 1999), and flies (Mamiya et al., 2018). FeCO neurons are selectively active over a certain range of positions, velocities, or vibration frequencies. Thus, they are range-fractionated, which allows the collective organ to expand its sensory range without compromising its sensitivity (Mamiya et al., 2018; Matheson, 1992b). FeCO is thus a multi-functional organ with functionally specialized sensilla.

FeCO afferents to the central nervous system (CNS) modulate the activity of leg motor neurons in a context-dependent manner. In an inactive insect, externally imposed movements of the tibia elicit a resistance reflex which is mediated by negative feedback from the FeCO (Agrawal et al., 2020; Field & Matheson, 1998). In active insects on the other hand, a positive feedback reinforces the ongoing movement of the tibia during the swing phase of walking (U. Bässler, 1976, 1988). Apart from this local proprioceptive role, FeCO is involved in body-wide responses to external stimuli both as a proprioceptor and as a vibroceptor. In crickets, FeCO mediates the induction and maintenance of thanatosis (Nishino & Sakai, 1996; H. Nishino et al., 1999). In fruit flies, FeCO club neurons detect low amplitude vibrations of 100 to 2000 Hz (Mamiya et al., 2018) and ablation of FeCO abolishes the freezing response of females to substrate borne vibrations produced by males during courtship (McKelvey et al., 2021).

Understanding the function of FeCO presents an intriguing challenge. Whereas FeCO is versatile in its functionality, reflecting the behavioural diversity of insects, its basic structure and mode of action are likely to be conserved across species (Field & Matheson, 1998; Virdi & Sane, n.d.). Moreover, both the neurobiology and mechanics are essential components of the mechanosensory function of the FeCO (Sane & McHenry, 2009). This makes FeCO ideal for comparative analyses, potentially providing insights into how evolutionary forces shape the structure and function of a mechanosensory organ. Our knowledge of FeCO morphology, anatomy and function is derived from studies in both hemimetabolous insects such as locusts, crickets, stick insects (Field & Matheson, 1998) and heelwalkers (Eberhard et al., 2010), and from holometabolous insects such as beetles (Frantsevich et al., 2019; Takanashi et al., 2016), flies (Mamiya et al., 2018, 2023; Shanbhag et al., 1992), true bugs (Hiroshi Nishino et al., 2016) and moths (Consoulas et al., 2000).

Here, we present a detailed anatomical description of FeCO in the Oleander hawkmoth *Daphnis nerii*. We used X-ray microtomography (microCT) to characterize the organ’s location inside the femur. To reveal the internal organization of FeCO, we used fluorescent staining and confocal microscopy, which allowed us to estimate the number of neurons and describe the morphology of actin-rich scolopales. We also traced the sensory neurons to visualize their central projections in the ventral nerve chord. Together, these data provide a comprehensive description of the FeCO anatomy in hawkmoths, as a first step towards characterizing their overall function.

## 2. Methods

### 2.1 Moth Culture

*D. nerii* moths were obtained from our laboratory culture at NCBS, Bengaluru, India. We reared the 1st to 5th instar larvae on *Nerium oleander* leaves. The pupae were kept in sawdust boxes. Adult moths were used within 1-3 days post eclosion. Males and females were released into a mating cage with plants for egg-laying and sugar solution for feeding. Eggs were collected from *N. oleander* leaves inside the cage and allowed to hatch in a container. We routinely introduced wild-caught larvae, pupae and moths into our lab culture to prevent inbreeding.

### 2.2 X-ray microtomography (MicroCT) imaging of femur

For the microCT scans, we enhanced the contrast in the soft tissue of femur by staining the samples with phospho-tungstic acid (PTA) using a protocol that we adapted for the purpose of imaging FeCO, as outlined below:

First, the legs were excised at the trochanter-femur joint and the distal half of tibia was cut away. The excised legs were permeabilized for 2-3 hours in phosphate-buffered saline with 0.3-0.5% Triton-X (PBST). The process of permeabilization involves treatment with a detergent which allows water to interact better with the hydrophobic cuticle, allowing stains and dyes to diffuse through to the tissue. After permeabilization, the legs were fixed with Bouin’s fixative for 5-6 days at 4°C. Excess fixative was removed by washing the samples with distilled water 3 times, 1 hour each wash. Before transferring the samples to staining solution, they were re-permeabilized in PBST for up to 3 days. To visualize the scoloparia, we stained the femur with a solution of 0.5% PTA in distilled water for 10 days. Because PTA stains cuticle lightly and has low penetration in long structures such as the femur, we altered the staining protocol on a subset of samples. We pre-stained the legs with aqueous iodine potassium iodide (IKI) to enhance the contrast of the cuticle for up to a month followed by incubation in 0.5% PTA solution with 10% methanol for several months to enhance penetration into the femur. Excess stain was washed off with distilled water 3 times, 30 minutes each wash. The samples were mounted in Vaseline® in a polypropylene micropipette tip and imaged in a micro-CT scanner (Bruker Skyscan 1172, Kontich, Belgium). The images were reconstructed using Bruker NRecon software (Bruker, Kontich, Belgium), and the femoral muscles, leg nerve and FeCO were segmented using Amira (version 5.4.3, FEI Visualization Sciences Group, France and Zuse Institute, Germany).

### 2.3 Fluorescent dye labelling of sensory soma and scolopale cells

To visualize and count the number of neurons in the FeCO in *D. nerii* and to gain insights into their organization into sensilla, we stained the organ using various dyes and imaged it using confocal microscopy. We stained the neurons, dendrites and caps using Neurobiotin 488 tracer (Vector Labs, USA). This method to obtain bright staining of dendrites and caps using neurobiotin is novel, and very effective for the visualization scolopidial morphology. We used phalloidin to visualize the scolopales which are composed of actin (Wolfrum, 1990). First, all moth legs were severed at the trochanter-femur joint. Breaking this joint allowed us to stain the FeCO in the proximal to distal direction *via* the proximal opening in the femur. The proximal end of the femur was first dipped in phosphate-buffered saline with 0.5% Triton-X (PBST) for 2-3 seconds and immediately transferred to a Vaseline® well of neurobiotin 488 tracer (Vector Labs, USA) solution prepared in 1M NaCl. The stain was allowed to diffuse into the femur for 45 minutes at 25°C. The legs were removed from the dye solution and fixed in 4% Paraformaldehyde (PFA) for 2 hours at 25°C. After washing off excess PFA, the legs were either mounted whole after removing a window in the ventral cuticle, or dissected to isolate FeCO. To isolate FeCO from femur, the FeCO apodeme was cut close to the joint and the proximal end of FeCO was cut free of its attachment to the femoral cuticle, near the Trochanter-Femur joint. The whole femur or the isolated FeCO was permeabilized in PBST (0.5% Tri-X) for 10 minutes and then stained with phalloidin conjugated with either Alexa Fluor 568 or Alexa Fluor 647 (Invitrogen, Thermo-Fisher Scientific, Massachusetts, USA), at 1:500 dilution in PBST (0.5% Tri-X) for 1-2 hours. After 2-3 washes in PBST (10 minutes each), the whole femur or the isolated organ was mounted on a slide in VectaShield. Both preparations were imaged using a confocal laser scanning microscope (Olympus FV3000; Shinjuku, Japan). Z-projections and 3D stacks were created using FiJi (ImageJ; National Institutes of Health).

### 2.4 Comparison of scolopale sizes

To determine the relative scolopale sizes in the two scoloparia, we used the confocal images described in the previous section to measure the scolopale sizes in 3D. We segmented each scolopale in 3D as a separate segment and extracted their Ferret diameters (diameter of a sphere that perfectly fits the segment) using the segment statistics module in 3D Slicer (Fedorov et al., 2012). Because scolopales are oblong, the ferret diameters quantify their length, regardless of their orientation within the tissue. The number of scolopales in dFeCO is an order of magnitude higher than its number in the vFeCO, we therefore used the non-parametric Wilcoxon Rank-sum test for statistical analysis of the scolopales in these two scoloparia.

### 2.5 Retrograde tracing of FeCO neurons

To visualize the axonal projection patterns of the FeCO sensory neurons in the CNS, we anterogradely filled the sensory cells. The moth was first immobilized ventral side up with dental wax and legs were descaled using a brush. The FeCO, which is superficially located, was exposed in the femur by peeling away the cuticle with a scalpel blade. The exposed FeCO was pricked by a sharp needle at random locations, and Dextran, Texas Red (Invitrogen, Thermo-Fisher Scientific, Massachusetts, USA, 3000 MW, lysine fixable, emission at 615 nm, excitation at 595 nm) diluted in very little water was applied to the organ. The wound was either covered with Vaseline® or allowed to clot. The moth was fed with sugar solution and kept alive for 30-32 hours for prothoracic FeCO and 20-24 hours for meso– and meta-thoracic FeCO fills. After incubation, the abdomen was cut away and the head and thorax were immersed in 4% PFA for overnight fixation at room temperature. The CNS was dissected out in PBS (pH 7.4) and dehydrated with the following ethanol grades: 50%, 60%, 75%, 90%, 100%, 100%. Samples were cleared and mounted in methyl salicylate between 2 coverslips and a metallic spacer for visualization in the confocal microscope (Olympus FV3000; Shinjuku, Japan) The acquired XYZ stacks were processed further in FiJi (ImageJ; National Institutes of Health) to enhance contrast and produce Z projections for the XY view and X projections for the YZ view.

## 3. Results

### 3.1 Location of FeCO

The FeCO is located in the proximal femur, with its distal end attached to a tanned apodeme which in turn connects to the tibia (Fig 1A, B). The X-ray microtomography images, when segmented, enabled identification of FeCO and its accessory structures within the femur. The femur houses 3 muscles; the antagonistic tibial flexor and extensor muscles that flex and extend the tibia respectively, and the retractor unguis (Fig 1B, C), a small muscle that moves the tarsus (Radnikow & Bassler, 1991; Snodgrass, 1935). The large extensor and flexor muscles are located opposite each other and are separated by a large space filled with trachea and haemolymph. FeCO is located in the ventral half of this space, closer to the ventral cuticle (arrow, Fig 1Ci) (Movie S1). The retractor unguis is located close to the dorsal cuticle in between the extensor and flexor and runs close to the leg nerve (Fig 1Ci-ii). In the medial femur, the FeCO apodeme moves away from the ventral cuticle towards the centre (arrowhead Fig 1C iii-iv). Closer to the femoro-tibial joint, the cross-sections of the muscles become smaller and the FeCO apodeme moves to a medial location close to the extensor muscle (arrowhead, Fig 1Cv-vi). The apodeme, which is chitinous, has a smaller cross-sectional area but greater stiffness than the FeCO, which is composed of soft neural and connective tissue, implying that FeCO and its apodeme operate as two elastic elements linked in series.

**Figure 1.**
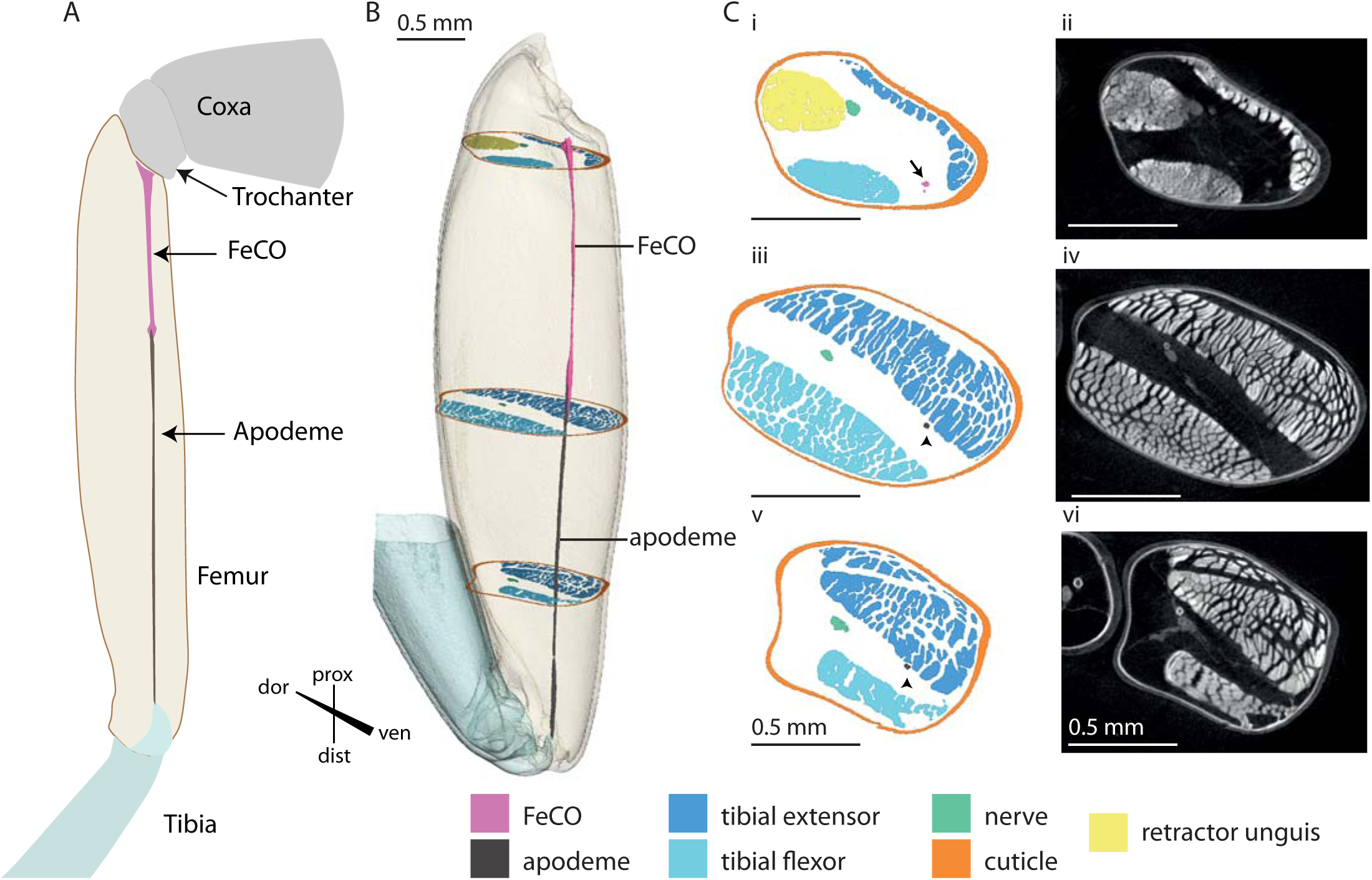
X-Ray Microtomography reconstruction of the FeCO and FeCO apodeme within the moth leg. **A.** Schematic shows the location of FeCO within the femur. Proximally, the FeCO is connected to the cuticle near the trochanter-femur joint. Distally, it attaches to a tanned FeCO apodeme, which in turn attaches to the tibia. **B. i.** 3D reconstruction of X-Ray Microtomography images show the location of FeCO (magenta) and FeCO apodeme (dark brow n) within the femur. **C.** Cross-sections of the segmented **(i, iii, v)** and unsegmented 3D reconstructed femur **(ii, iv, vi)** at a proximal location **(i, ii)**, medial location **(iii, iv)**, and distal location **(v, vi)** show the location of FeCO (magenta, arrow) and FeCO apodeme (dark brown, arrowheads) relative to the extensor muscle (dark blue), flexor muscle (light blue), retractor unguis muscle (yellow) and the leg nerve (green). All scale bars are 0.5 mm. prox= proximal, dor= dorsal, dist=distal and ven=ventral.

### 3.2 Composition of FeCO

The FeCO is divided into two distinct scoloparia which are visible at its proximal end (Fig 2Ai-ii). Based on their location, they may be defined as *dorsal* (dFeCO) and *ventral* (vFeCO) scoloparia. Of the two scoloparia, dFeCO is larger than vFeCO. Both scoloparia are wider at their proximal ends and taper towards their distal end. Approximately halfway along the length of the organ, they fuse and continue distally to attach to the apodeme (Fig 2Aii, Bi). They are thus connected in parallel between their proximal and distal attachment points. vFeCO is closer to the ventral cuticle of femur (Fig 2B ii), whereas dFeCO is located deeper within the femur (Fig 2B iii).

**Figure 2.**
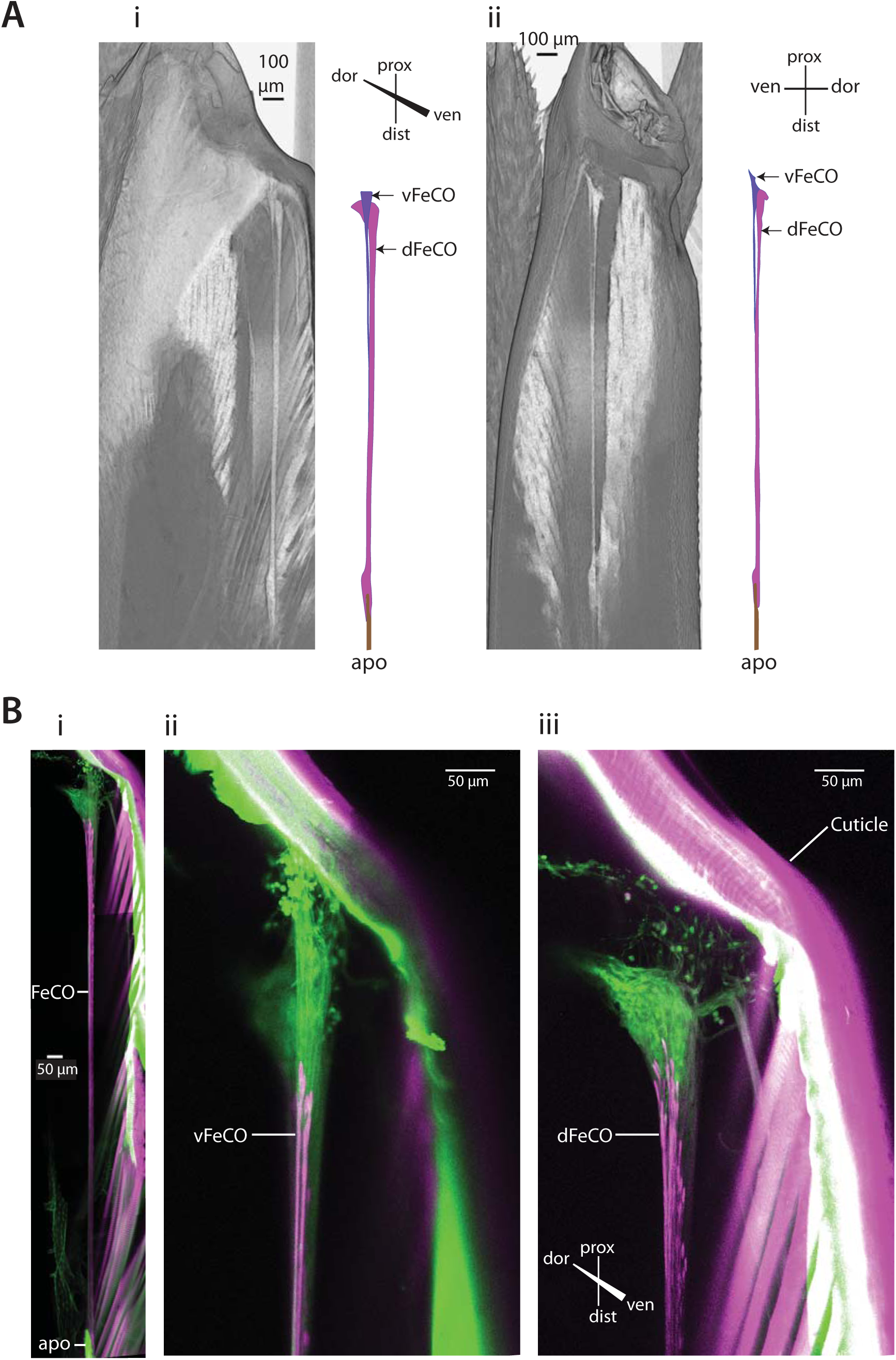
The ventral and dorsal scoloparia of the FeCO in *D. nerii*. **A.** Dorsal **(i)** and lateral **(ii)** views of virtually sliced 3D reconstructed femur showing the location and arrangement of FeCO scoloparia. The ventral scoloparia (vFeCO) lies just underneath the ventral cuticle whereas the dorsal scoloparia (dFeCO) lies deeper within femur. Distally, it connects to the FeCO apodeme (apo). Scale bars=100 mm **B.** Confocal z-stack of FeCO in a whole mount of femur showing its connection to femoral cuticle proximally and FeCO apodeme distally **ii.** Z-stack showing vFeCO with a faint shadow of the dFeCO behind it. **iii.** Z-stack showing dFeCO which lies behind vFeCO (not visible). Scale bars=50 mm

### 3.3 Organization and soma size variation in FeCO

To visualize the internal organization of FeCO in moths, the neurons were filled with neurobiotin, and the actin fibres of the scolopales were stained with phalloidin. Confocal imaging of a whole mount of femur revealed the proximal location of sensory soma within FeCO (Fig 2Bi). We separately imaged an isolated FeCO to obtain high resolution scans showing individual sensory neurons, dendrites, caps and actin fibres (Fig 3). In addition to marking the actin-rich scolopales, phalloidin staining also revealed the presence of longitudinally oriented actin filament along the entire length of the FeCO ligament (Fig 3A). Cross-sections of FeCO at the attachment cell level (distal to scolopales) (Fig 3B i-iii) show the ring-like concentration of actin fibres (magenta) at the periphery of attachment cells, contained within a thin sheath of neural lamella (green) (Movie S2). Here, the neural lamella fluoresced in the green spectrum either due to the non-selective staining by neurobiotin or due to autofluorescence. The unstained attachment cells appeared as hollow spaces at the center of the actin rings and neural lamella.

**Figure 3.**
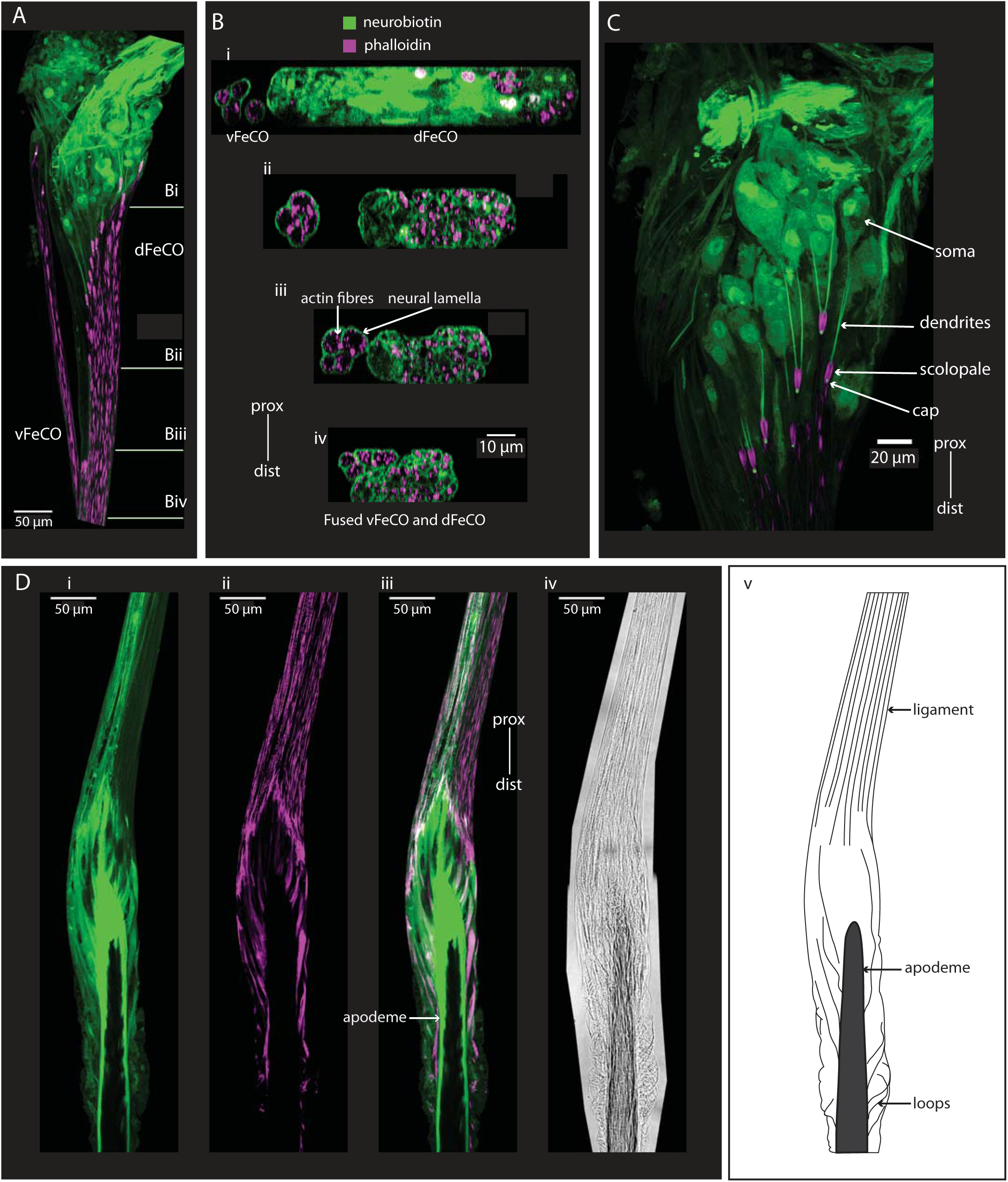
Organization of neurons, scolopale and actin fibres in the FeCO. **A.** Imaging of isolated FeCO allows for a scan of higher resolution showing the internal organization of sensory soma and scolopales. **B. i-iv** Cross-sectional views of **A** showing a neural lamella stained with neurobiotin covering each attachment cell of the vFeCO, containing actin fibres close to the periphery. The 4 attachment cells of the vFeCO can be traced distally (i-iii) until they merge with the dFeCO (iv) **C.** A Z-projected confocal image showing 2 dendrites projecting to a single scolopale (magenta) as they terminate in the neurobiotin stained caps (green), surrounded by the scolopale. **D.** Individual fibres in FeCO ligament connect to the apodeme at various points. **i.** Neural lamella in the FeCO ligament stained with neurobiotin (green). **ii.** Actin filaments in the distal part of the FeCO ligament stained with phalloidin (magenta). **iii**. merged image. **iv**. Distal-most fibres attach to the sides of the apodeme and show loop like appearance. **v.** Cartoon diagram of the ligament and its connections to the apodeme. Scale bars=50 mm

In the proximal part of FeCO, the larger and proximally-located soma were more circular whereas the smaller distally-located soma were slender and long (green, Fig 3C). The dendrites of FeCO sensory neurons (green, Fig 3C) project distally towards the tibia and are enveloped by the actin rods of a scolopale (magenta, Fig 3C). Because neurobiotin did not stain all sensory neurons, we used the phalloidin-stained scolopales to count the number of sensory neurons. The small vFeCO is made up of 8 neurons (4 scolopales) and the larger dFeCO has 94-108 neurons (47-54 scolopales).

The attachment cells extended distally to join the tongue-shaped tip of the apodeme at various locations (Fig 3D). Some cells connected to the tip of the tongue while the others ran parallel to the tongue before joining it on the side. We observed some looping in the attachment cells that join the apodeme most distally (Fig 3D i, iii). The actin filaments were present in the entire length of the attachment cells (Fig 3D ii-iii). The attachment cells were clearly visible as fibres under transmission lighting (Fig 3D iv).

### 3.4 Scolopidial units and scolopale sizes

The dendritic terminals, surrounded by an actin-rich scolopale, terminated in the cap which in turn connected to the attachment cell (Fig 4A, B). The dendrites contain microtubules whereas the caps are made of an electron-dense material secreted from scolopale cells (Yack & Roots, 1992) or the attachment cell (Moran et al., 1975). We quantified the scolopale sizes of the vFeCO (Fig 4C i, ii) and dFeCO (Fig 4C iii, iv) by segmenting them (Fig 4D i) and measuring their feret diameter, i.e., diameters of spheres that completely enclosed individual 3D segmented scolopales (3D Slicer) (Fig 4D ii). As evident from these measurements (Fig 4E), vFeCO scolopales sizes are substantially larger than dFeCO scolopales (Wilcoxon Rank-Sum test, p < 0.0001). The scolopale sizes in the vFeCO lie in the range of 17.4 – 23.5 μm (mean: 20.3 μm, SD: ±1.25 μm), whereas in the dorsal scoloparia, scolopale sizes are more tightly distributed between 9.7 – 16.3 μm (mean: = 13.3 μm, SD: ± 1.11 μm). As previously noted, the scolopale numbers are also very different; the dorsal scoloparia contains 47-54 scolopales, whereas the ventral scoloparia contains only 4 scolopales.

**Figure 4.**
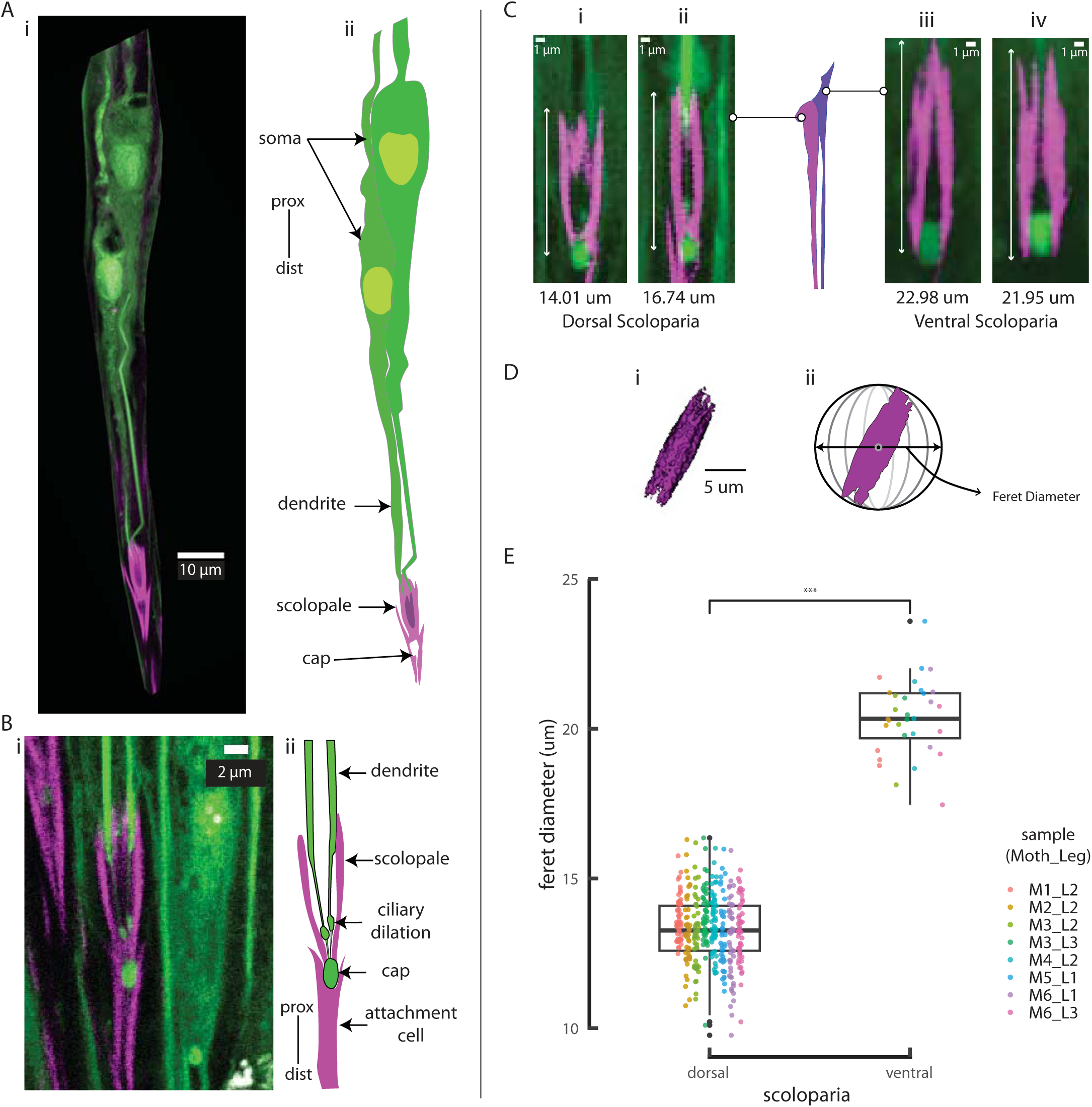
Relative sizes of scolopales in the vFeCO and dFeCO. **A.** Confocal image (left) and traced diagram (right) of a single scolopidial unit showing (from top to bottom) sensory neuron soma, sensory dendrite, scolopale cell, cap. Scale bar=10 mm **B.** confocal image (left) and traced diagram (right) of a scolopale (magenta) surrounding 2 sensory dendrites (green) which show a ciliary dilation just before terminating in the cap (green). Distal to the caps, actin filaments stained with phalloidin (magenta) mark the edges of the attachment cells that join the scolopidial unit to the apodeme. Scale bar=2 mm **C.** Confocal images of 2 scolopales from ventral scoloparia and dorsal scoloparia showing the difference in their lengths. The lengths are shown under each figure. Scale bar=1 mm **D.** To measure the length of 3D segmented scolopales (**i**), we used the diameter of a sphere which circumscribes the 3D segmented scolopale (feret diameter) (**ii**). Scale bar= 5 mm **E.** Box plots showing the significant different (Wilcoxon Rank-Sum Test, p < 000.1) between scolopale sizes from the ventral and dorsal scoloparia. The samples are identified using labels “MX_LY”, where X depicts the sample number, and Y depicts the leg identity (1=prothoracic, 2=mesothoracic and 3=metathoracic). Each sample is depicted by a single color, and each data point in the box plot represents the feret diameter value of a single scolopale within that sample.

### 3.5 FeCO neurons project to the ipsilateral thoracic ganglia

To trace the central projection patterns of the neurons within the FeCO, we labeled them with fluorescent dyes. The axons of the FeCO from each leg project to their respective hemiganglion. For example, prothoracic FeCO neurons enter the prothoracic ganglia *via* either the foreleg nerve IN1b or IN2a (Eaton, 1974), before terminating in the medio-posterior region of the ganglia (Fig 5A i-ii). These arbors are restricted to the ventral half of the ganglia and terminate close to the midline (Fig 5A iii). In contrast, the mesothoracic (Fig 5B) and metathoracic FeCO afferents (Fig 5C) enter their respective ganglia at the posterior end through nerve IIN5 and IIIN2 (Eaton, 1974) respectively and project medio-anteriorly (Fig 5B i-ii, C i-ii). Like the prothoracic FeCO projections, their arbors are restricted to the ventral side of the ganglia and again terminate very close to the midline (Fig 5B iii, C iii). Mesothoracic FeCO projections terminate near the Ventral Intermediate Tract (VIT) (Suder & Wendler, 1993) (Movie S3). The leg motor neurons in all three ganglia, as well as the flight motor neurons in the meso– and metathoracic ganglia are restricted to the dorsal half of the ganglia. Because we dye-filled the neurons by injuring them sparsely with a sharp needle, only a subset of neurons got stained in each preparation and therefore the full central projections of all FeCO neurons could not be visualized at once.

**Figure 5.**
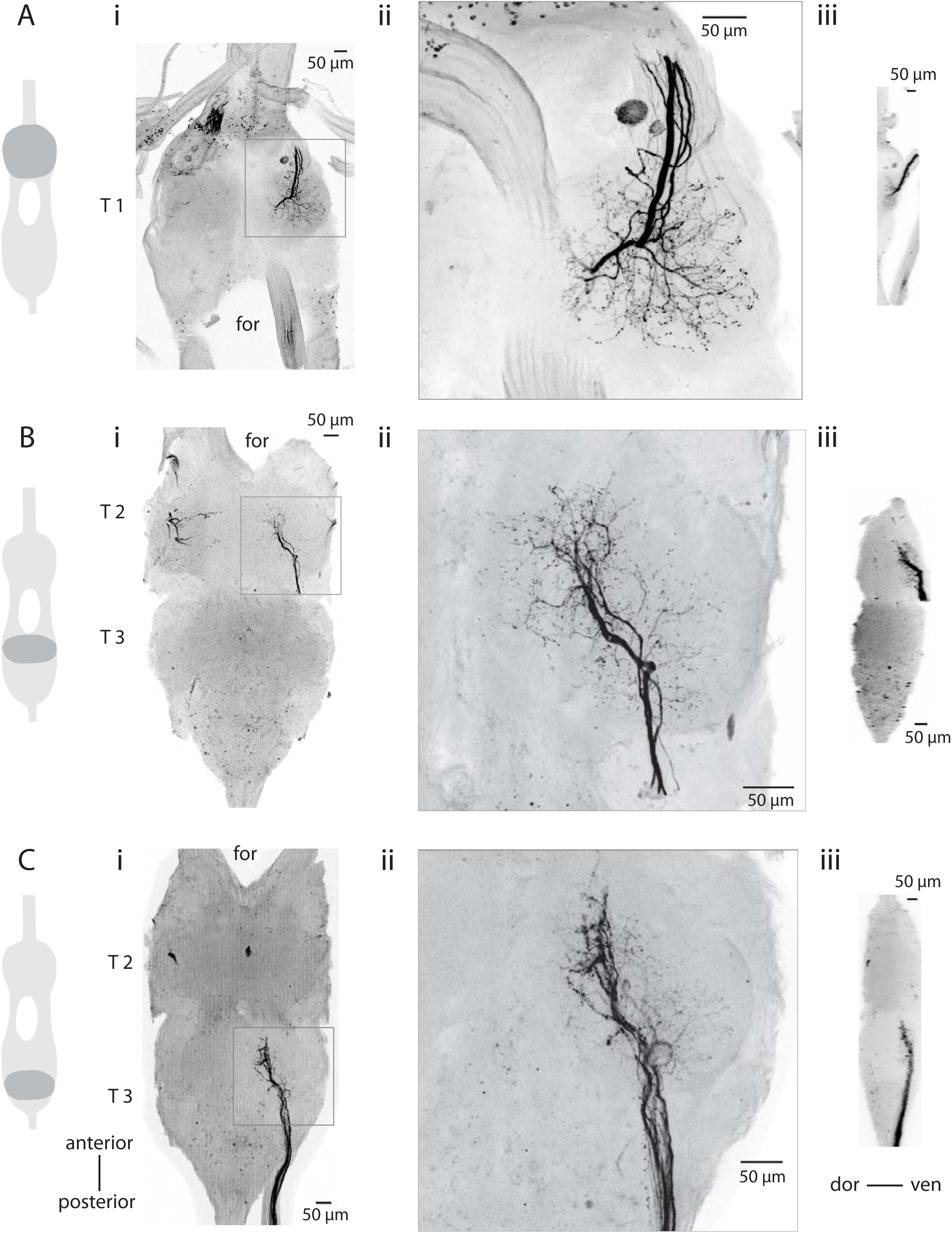
Central projections of FeCO sensory neurons. Central projections of the FeCO projections in the prothoracic ganglion (T1) (Ai-iii), mesothoracic ganglion (T2) (Bi-iii) and metathoracic ganglion (T3) (Ci-iii). In each subfigure, the leftmost column shows the schematic of the part of the ventral ganglion that was imaged, (i) depicts a zoomed-out image of the projections, (ii) depicts the enlarged image (indicated by the box in (i), and (iii) depicts the laterally re-sliced and z-projected view. All scale bars =50 mm.

## 4. Discussion

In this study, we conducted a detailed anatomical and morphological investigation of FeCO in the Oleander hawkmoth *D. nerii*. First, we developed a novel microCT staining protocol with aqueous PTA to image and reconstruct an intact femur in 3D, which enabled us to observe the locations of muscles and nerves relative to FeCO. In addition, we pre-stained the leg samples with IKI for better staining of the cuticle, which allowed us to segment the femoral and tibial cuticle. Second, we examined the internal structure and composition of the hawkmoth FeCO by staining it with neurobiotin which revealed the relative size and organization of the underlying neurons within the large (dFeCO) and small (vFeCO) scoloparia. We observed and characterized in detail the size differences in the scolopales of these two scoloparia using phalloidin. Third, we mapped the central arbors to determine the specific regions within the thoracic ganglia where the sensory signals from the FeCO are transmitted. This study demonstrated that FeCO arbors terminate locally within their nearest associated ganglion approaching but not crossing the midline. Together, these data lay the basis for a broader understanding of the FeCO structure and function from a physiological and a comparative standpoint, as discussed in the following sections.

### 4.1 A mechanical model of the FeCO structure and function

In *D. nerii*, FeCO is located in the proximal femur and connected to the tibia *via* an apodeme (Fig 6A). The compliant FeCO tissue undergoes stretch and relaxation during cyclic activity. If the apodeme is compliant, then depending on the elastic modulus of the apodeme, part of the stretch may be absorbed, effectively dampening the strain transferred to the FeCO tissue. Alternatively, an inextensible apodeme may behave as a strut and transfer the stimulus without dampening it. Thus, measurements of the material properties of the FeCO apodeme are essential for determining the mechanical pre-filtering of stimulus.

**Figure 6.**
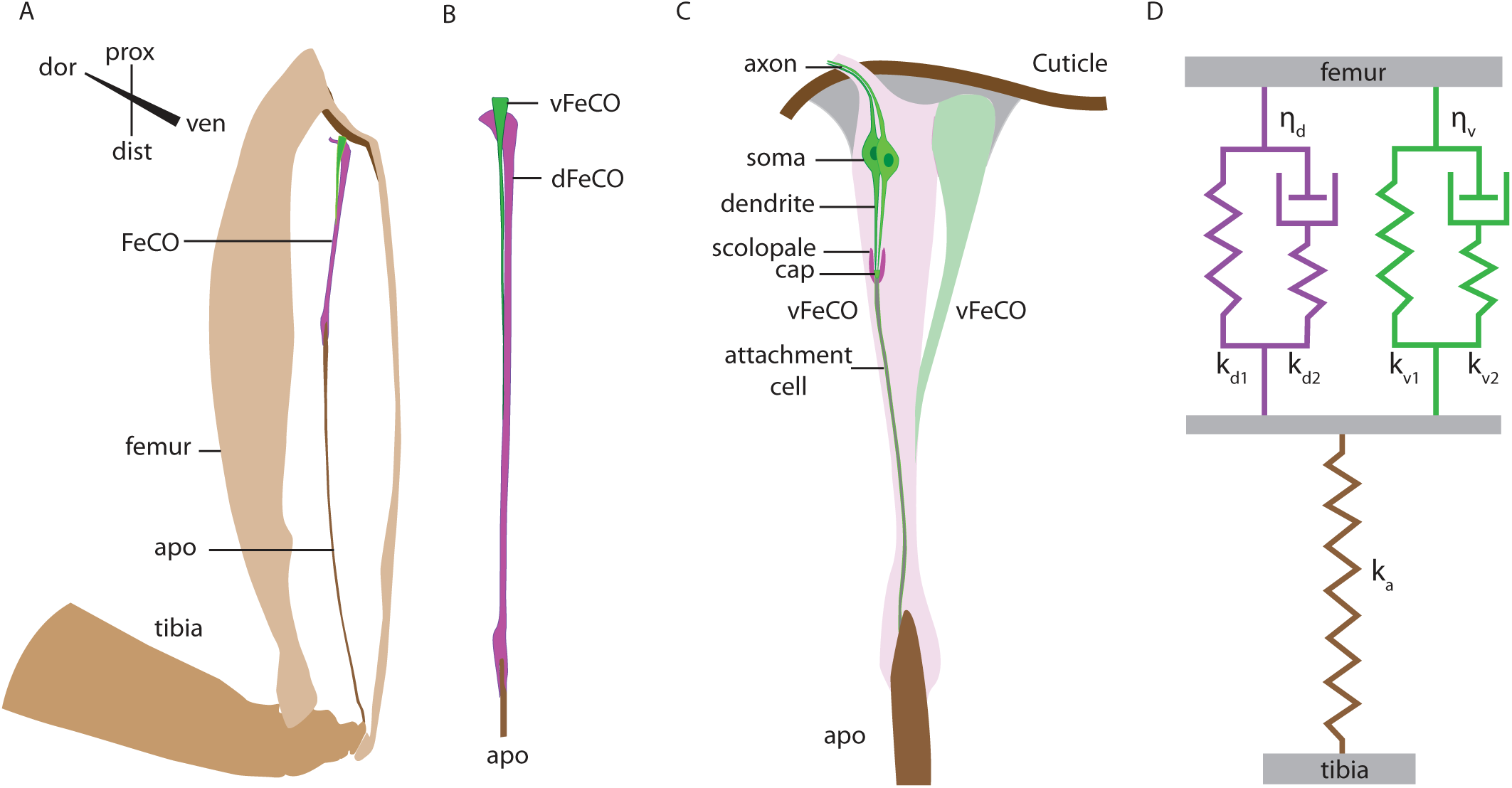
A mechanical model of FeCO in *D. nerii*. **A.** Schematic of FeCO in *D. nerii* shows its location in the proximal femur and connection to tibia via the apodeme. **B.** The two scoloparia of FeCO are differ in their sizes and connect to the apodeme in parallel, at their distal ends. **C.** Neurons in FeCO are organized into scolopidial units. Each unit consists of 2 sensory neurons, a scolopale, a cap and an attachment cell that connects directly to the apodeme. Scolopidial units within each scoloparium vary with respect to their soma size, scolopale size, point of attachment between the attachment cell and the apodeme, and their collective position within the scoloparia. These factors may influence their range selectivity. **D.** The difference in the geometries of the two scoloparia mean that they likely experience different stress for the same strain. Thus, they are modelled as two standard linear solid viscoelastic elements with different spring constants (k_v1_, k_v2_ and k_d1_, k_d2_) and viscosity constants (η_v_ and η_d_), connected in parallel to the apodeme (spring with constant k_a_).

The FeCO of *D. nerii* consists of two scoloparia, dFeCO and vFeCO which may be approximated as two cones connected in parallel to the apodeme, in which vFeCO is smaller than dFeCO (Fig 6B). These components are made of softer tissue, within which the mechanosensory neurons are embedded and sense the local deformation (Fig 6C). Moreover, previous electrophysiological studies have shown that the FeCO responses to cyclic loading and relaxation shows some hysteresis (Büschges, 1994; Matheson, 1990; Zill, 1985). FeCO also shows mechanical hysteresis (Walker, 1998), which may reflect in its physiological responses. Additionally, FeCO shows adaptation in its tonic responses when held under a constant FT joint position (Burns, 1974; Usherwood et al., 1968) and in its internal stress when held under a constant stretch (Walker, 1998), which also suggests that some material relaxation may occur with the FeCO tissues. Both hysteresis and adaptation are indicative of viscoelastic properties.

Together, these observations can be integrated into a mechanical model of the FeCO structure (Fig 6D). In this model, the dFeCO and vFeCO, which differ in cross-sectional areas, are represented as two viscoelastic elements. Each element consists of a spring in parallel with a spring and dashpot in series, following the standard linear solid model (Vincent, 1990). The two elements are arranged in parallel to represent the FeCO tissue, including dFeCO and vFeCO respectively. The FeCO tissue is then connected in series with an elastic element that represents the rigid apodeme (Fig 6D). For the uniaxial stress condition along the long axis, dFeCO and vFeCO would experience differential stresses (i.e. force experienced by the structure per unit cross sectional area) although they undergo the same strain (i.e. change in length experienced by the structure per unit length). Thus, this model presents a mechanical basis for range segregation of the mechanical signal experienced by the FeCO.

### 4.2 The composition and internal mechanics of FeCO

Each scoloparium consists of numerous parallelly arranged scolopidia comprising 2 neurons, 1 scolopale, 1 cap and 1 attachment cell, whose structure is conserved across insects (Fig 4A, 6C) (Field & Matheson, 1998; Virdi & Sane, n.d.). However, the scolopidia morphologically differ from each other in their neural soma size, scolopale size, point of connection between the attachment cell and the apodeme, and position within the scoloparia (Fig 6D). The cone-shaped geometry of the scoloparia suggests that the stress within each scoloparia is not homogeneously distributed, and hence the sensitivity of the soma within FeCO could be dependent on its location which determines its local stimulus field.

In the data presented here, soma sizes vary with the position of their scolopidial unit within the scoloparia. For instance, within the dFeCO, the larger round soma are located proximally, where the cross-sectional area is larger, whereas, the smaller slender soma are located distally, where the cross-sectional area is smaller. From a morphological viewpoint, these observations suggest range selection of the neurons dictated by their location within the organ. For instance, in the vinegar fly FeCO, the position sensitive claw neurons are activated by tibial flexion in a sequential manner from the most proximal to most distal. Their activity thus depends on their location within the scoloparia as the proximal region gets stretched first, followed by the distal region (Mamiya et al., 2023). Intracellular recordings from FeCO afferents accompanied with dye-fills can help generate a functional map of FeCO neurons within the organ. Such structural features may additionally contribute to range fractionation of the scolopidial units in the FeCO (Fig 6D).

The scolopale sizes differ significantly between the two scoloparia, with an average length of 20.3 μm in vFeCO and 13.3 μm in dFeCO (Fig 4). Whether this size difference has physiological consequences remains an open question, requiring selective ablation or recordings. Additionally, measuring the scolopale sizes in FeCO scoloparia with known functional specialization will help establish the relationship between scolopale size and scoloparia function. Although scolopale sizes are shown to vary in the auditory chordotonal organ in the parasitoid fly *Ormia ochracea* (Robert & Willi, 2000), such size characterization is lacking in the FeCO of other insects.

The attachment cells that form the ligament of the organ connect to the apodeme in parallel *albeit* at spatially separated points along its tongue-shaped tip (Fig 3D). In the metathoracic FeCO of locusts, the ligament attaches to the apodeme at various points and appears as coiled loops at the extended position which suggests that they are not under tension. When the apodeme is displaced distally upon tibial flexion, different loops experience tension at different displacements of the apodeme (Field, 1991). Similarly, in the cricket FeCO, the apodeme extends into the FeCO ligament as a coiled core to which attachment cells connect at various points. When the apodeme displaces distally during tibial flexion, the coiled cores unfold and straighten out, likely bringing the attachment cells under tension one by one (Nowel et al., 1995), causing sequential recruitment of individual scolopidial units.

### 4.3 The role of actin filaments in FeCO mechanics

Phalloidin stains revealed actin fibres arranged in a ring-like shape at the periphery of attachment cell cytoplasm (Fig 4 Bi-iv), which is previously unreported. Attachment cells are connected to each other *via* cell-cell junctions (Lipovsek et al., 1999) and are covered by the connective tissue sheath, the neural lamella, which wraps individually around each attachment cell (Field, 1991; Young & Pringle, 1970). Actin filaments may form a part of the cytoskeletal support at the cell-cell junctions or provide elasticity to the FeCO ligament as contractile actin.

### 4.4 Comparison of FeCO central projections across insects

FeCO central projections have not been previously described in moths. Consistent with the central projections in locusts (Field & Pflüger, 1989), stick insects (Schmitz et al., 1991), and crickets (Nishino & Sakai, 1997), *D. nerii* FeCO neurons from each leg project ipsilaterally and terminate within their respective hemiganglia. These projections are confined to the ventral half of the thoracic ganglia (Fig 5). In *D. nerii* mesothoracic FeCO neurons terminate near the Ventral Intermediate Tract (VIT). In many insects, the two scoloparia show distinct central projection patterns; whereas one scoloparia sends projections to mVAC, the other terminates in lateral association centres (LACs). For instance, in locusts, neurons from the proximal scoloparia arborize in LAC whereas the distal scoloparia afferents terminate in the medial Ventral Association Centre (mVAC) (Field & Pflüger, 1989). Similarly in crickets, the ventral scoloparia neurons terminate in the dorsolateral region between the VIT and the dorsal intermediate tract (DIT) and at the periphery of mVAC whereas the dorsal scoloparium sends neurons exclusively to the mVAC (Nishino & Sakai, 1997). Central projections in moths may similarly follow distinct patterns for the two scoloparia, but their separate staining presents some difficulties. Unlike in locusts pro– and meso-thoracic legs the nerves of the proximal and distal scoloparia of FeCO are individually accessible, staining vFeCO and dFeCO is more difficult as they are very closely adjoining.

In locust FeCO, soma of physiologically similar neurons are grouped together (Matheson, 1990), with some correlation between sensory responses and central projection patterns (Matheson, 1992a). In vinegar fly, position sensitive claw neurons show goniotopic (based on joint angle) organization in the organ and the vibration sensitive club neurons show a tonotopic (based on vibrational frequency) organization of their dendrites within FeCO (Mamiya et al., 2023) as well as in their axons in the thoracic ganglia (Mamiya et al., 2018). Electrophysiological measurements combined with dye-filling are essential to ascertain whether FeCO neurons in *D. nerii* neurons are topologically organized both in the CNS and within their respective FeCO scoloparia.

The localization of FeCO projections to their respective hemiganglion suggests that they provide local sensory feedback to the leg motor system. In locusts, FeCO neurons make monosynaptic reflex arcs with the flexor tibiae motor neurons (Burrows, 1987) which allows for rapid flexor motor responses to externally imposed extension stimuli (reviewed in Field & Matheson, 1998). Afferents also connect to both spiking (Burrows, 1987, 1988), non-spiking (Burrows et al., 1988), and intersegmental interneurons (Laurent & Burrows, 1988). Similarly in stick insects, FeCO neurons connect to spiking (Ansgar Büschges, 1989) and non-spiking interneurons (Ansgar Büschges, 1990). Parallel connections to motor neurons and various interneurons mediate the afferent control of the leg motor system.

### 4.5 Comparison of the FeCO anatomy in moths with other insects

The proximal location of FeCO in *D. nerii* (Fig 6A) is consistent with FeCO in the lepidopteran tobacco hornworm moth (Consoulas et al., 2000; Kent & Griffin, 1990), dipteran vinegar fly (Shanbhag et al., 1992) and blowfly (Lakes & Pollack, 1990), orthopteran locusts (Eleanor H. Slifer, 1935), crickets (H. Nishino & Sakai, 1997), and New Zealand weta (Hiroshi Nishino, 2003), and phasmatoid stick insects (Hofmann et al., 1985). Conversely, in coleopteran Japanese pine sawyer beetle (Takanashi et al., 2016), neuropteran lacewings (Lipovsek et al., 1999) and hemipteran cicadas and stink bugs (Hiroshi Nishino et al., 2016), FeCO is situated in the medial femur, whereas in the specialized hindlegs of locusts, it is located close to the FT joint (Usherwood et al., 1968). Although FeCO location and gross morphology varies across insects, the scolopidial structure of FeCO in *D. nerii* is conserved across locusts (Moran et al., 1975; E. H. Slifer & Sekhon, 1975), flies (Shanbhag et al., 1992) and lacewings (Lipovsek et al., 1999).

In *D. nerii*, whereas the dFeCO comprises approximately 94-108 neurons, the vFeCO is an order of magnitude smaller with only 8 neurons. This is consistent with data from the tobacco hornworm moth *Manduca sexta,* also from the same Sphingidae family, which has a large scoloparium (nFeCO) composed of 45-60 neurons that grows *de novo* in the larval-to-pupal stage of development, and a smaller scoloparium (lFeCO) that has 6 neurons and is larval-persistent (Consoulas et al., 2000). Similarly, in locusts the proximal and distal scoloparium contain approximately 200 and 50 neurons respectively (Burns, 1974) and in stick insects, dorsal scoloparium has 400 while the ventral has 80 neurons (Fuller & Ernst, 1973 as cited in Hofmann et al., 1985). Interestingly, the size difference correlates with their function as the proprioceptive scoloparia is many times smaller than the vibroceptive scoloparia in both locusts and stick insects (Field & Pflüger, 1989; Kittmann & Schmitz, 1992). It is not yet determined if the smaller vFeCO and larger dFeCO in *D. nerii* are similarly proprioceptive and vibroceptive respectively.

### 4.6 FeCO compared to other chordotonal organs

Like the morphology and functions of the legs in which they are housed, FeCO function may widely vary across insects. It has been proposed that the Johnston’s Organ (JO) in the insect antenna which detects passive flagellar movements and vibrations, may have evolved from the FeCO when antenna evolved from legs (Krishnan & Sane, 2015). For instance, studies in stick insects show that ablating the antenna between scape and pedicel at the nymphal stages leads to development of the 2 distal most segments of the leg – tibia and tarsus in the next molt (Dürr, 2016). There are similarities in their underlying sensorimotor system and their mechanosensors, including homologies between the hair sensilla on the antennal segment scape to those on proximal leg segments (Krishnan & Sane, 2015). Thus, ablation of hair sensilla in antenna (or Böhm’s bristles) disrupts antennal positioning (Krishnan et al., 2012) just as ablation of leg hair plates disrupts walking in locusts, cockroaches and stick insects (Ulrich Bässler, 1993; Cruse et al., 1984; Wong & Pearson, 1976). Moreover, both FeCO (Hofmann et al., 1985; Matheson, 1992b) and JO (Sane et al., 2007; Dieudonne, Daniel & Sane, 2014) exhibit the property of range fractionation, as determined through the measurement of their physiological responses. The hypothesis of homology between FeCO in legs and JO in antennae needs to be explored further to gain insights into the evolutionary origins of JO.

### 4.7 Conclusions

The diverse functional, morphological and physiological properties of FeCO make it an excellent model system to study the role of mechanical pre-filtering in mechanosensory physiology. Three hypotheses in particular emerge from the data presented in the current study. First, the two scoloparia of FeCO can be modelled as two viscoelastic elements parallelly connected to the apodeme, such that their individual spring and viscosity constants will determine the amount of stress experienced by the neurons they house, for the same amount of strain. Second, the morphological differences in the individual scolopidial units with respect to their scolopale sizes and position within the conically shaped scoloparia, and their point of attachment to the apodeme can determine their functional tuning to different ranges of FT joint position, speed and direction. In locusts and crickets flexion of tibia recruits individual sensillae sequentially (stretched) giving them a mechanical threshold for activation at different FT joint angles (Field, 1991; Nowel et al., 1995). Third, actin filaments in the FeCO ligament could contribute to its elasticity. Ultimately, electrophysiological studies of individual FeCO neurons, which also form the neural filter for the stimulus, would help us probe the hypotheses relating structure to function. Our current data lays the groundwork for future investigation into FeCO physiology and biomechanics.

## Supporting information

Virdi Sane Supplementary Movie 1

Virdi Sane Supplementary Movie 2

Virdi Sane Supplementary Movie 3

## Acknowledgements

We acknowledge the MicroCT facility at NCBS, TIFR, Bengaluru for the MicroCT scans. Mr. Sunil Prabhakar from the MicroCT facility helped develop the protocol for aqueous PTA staining with 10% methanol. We also acknowledge the Central Imaging and Flow Cytometry Facility (CIFF) at NCBS, TIFR for the confocal imaging. We acknowledge Mr. Kemparaju, Mr. Allan Joy, and Mr. Anil Kumar for managing the moth culture at NCBS, TIFR. Funding was provided by the Air Force Office of Scientific Research grants (AFOSR; FA2386-11-1-4057 and FA9550-16-1-0155) and NCBS, supported under project no. 12-R&DTFR-5.04-0900.

## Supplementary information legends

**Movie S1.** A 3D view of a reconstructed hindleg femur showing the location of FeCO and its apodeme, with and without segmentations of the femoral and tibial cuticle, extensor, flexor and retractor unguis muscles and the leg nerve. Cross-sections show the position of FeCO or its apodeme inside the femur at three separate locations; proximal, medial and distal femur.

**Movie S2.** A longitudinal and transverse views of isolated FeCO prep with neurons, dendrites and caps stained with neurobiotin (green), and filamentous actin stained with phalloidin (magenta). The transverse sections start from the proximal end (top) of the organ where the neural soma are located. More medially, scolopales appear, followed by ring-like arrangement of actin fibres at the periphery of attachment cells. The attachment cells appear hollow as they were not stained. A neural lamella, which was either stained with neurobiotin non-selectively, or auto-fluoresced in the green channel can be seen around each attachment cell individually.

**Movie S3.** A lateral view (stack) of the mesothoracic ganglia showing the central projections entering the ganglia from the leg nerve on the ventral (left) side. A movie showing slices in the ventral-dorsal axis, showing the extensive arborization of FeCO neurons in the leg neuropil as well as their termination near the Ventral Intermediate Tract (VIT).

